# Influence of a mono-frequency sound on bacteria can be a function of the sound-level

**DOI:** 10.1101/071746

**Authors:** Vijay Kothari, Chinmayi Joshi, Pooja Patel, Milan Mehta, Shashikant Dubey, Brijesh Mishra, Niral Sarvaiya

## Abstract

*Chromobacterium violaceum* was subjected to sonic stimulation with 300 Hz sound, at five different levels of loudness in the range of 70 – 89.5 dB. Sonic stimulation was found to affect bacterial growth and quorum sensing regulated pigment (violacein) production significantly. Magnitude of this effect was found to be dependent on sound-level. The minimum critical difference required to cause any statistically significant change in bacterial response with respect to sound-level was found to be 13 dB. Growth of *C. violaceum* was affected more at lower sound intensity, whereas pigment production was affected more at higher sound intensity. Additional experiments with *C. violaceum* and *Serratia marcescens* indicated that even a silent speaker emitting no sound can alter bacterial growth and/or pigment production upto a minor extent. Size of the test tube in which bacteria are exposed to sonic stimulation, was not found to affect the results much.

## Introduction

It is a common characteristic of all living entities to respond to external stimuli such as light, chemicals, etc. Microorganisms exert phototactic and chemotactic movement in response to light and chemicals respectively (Madigan *et al.*, 2009). Sound is one of the most ubiquitously present physical stimuli in the environment, to which almost all life forms are exposed to a varying degree. Despite sound being one of the most widespread environmental factors, not enough investigation has been there on how microorganisms sense and respond to the external sound stimulation. Such investigations can be considered an important aspect of the dynamics of microbial growth, wherein ‘dynamics’ refers to the study of those environmental forces which can either promote or impede microbial growth (Tempest *et al.*, 1978).

Influence of sound on higher animals having well-developed hearing organs can be understood relatively easily. There have been reports describing effect of sound (either in form of music or otherwise) on animals (Marten and Marler, 1977; Chanda and Levitin, 2013), plants (Sun *et al*., 1999), as well as, microorganisms (Aggio *et al.*, 2012; Shah *et al*., 2016). However, mechanisms of perceiving and responding to sound differ among these three life forms, as organisms other than animals do not posses specific auditory machinery. Our knowledge regarding how microorganisms interact with sonic range (20-20,000 Hz) sound is limited.

In the present study, we report our investigations regarding the effect of a mono-frequency (300 Hz) sound on the gram-negative bacterium *Chromobacterium violaceum* at different sound levels. Additionally, we also investigated whether the size of test tube in which bacteria are exposed to sound makes any difference to microbial response to external sonic stimulation.

## Materials and Methods

### Bacterial culture

*C. violaceum*(MTCC 2656) and *Serratia marcescens* (MTCC 97) was procured from the Microbial Type Culture Collection (MTCC), Chandigarh. Both the organisms were grown in nutrient broth (HiMedia, Mumbai) supplemented with 1% v/v glycerol, and incubated at 35° C and 28° C respectively.

### Sound generation

Sound beep(s) of required frequency was generated using NCH^®^ tone generator. The sound file played during the experiment was prepared by using WavePad Sound Editor Masters Edition v.5.5 in such a way that there is a time gap of one second between two consecutive beep sounds.

### Sound stimulation

Inoculum of the test bacterium was prepared from its activated culture, in sterile normal saline. Optical density of this inoculum was adjusted to 0.08-0.10 at 625 nm (Agilent Technologies Cary 60 UV-Visible spectrophotometer). The test tubes (Borosil, 25 x 100 mm) containing inoculated growth medium (total volume of the inside content was 6 mL including 5% v/v inoculum) were put into a glass chamber (Actira, L: 250 x W: 250 x H: 150 mm). A speaker (Minix sound bar, Maxtone Electronics Pvt. Ltd., Thane) was put in this glass chamber at the distance of 15 cm from the inoculated test tubes. Sound delivery from the speaker was provided throughout the period of incubation (48 h). This glass chamber was covered with a glass lid, and one layer of loose-fill shock absorber polystyrene, in such a way that the polystyrene layer gets placed below the glass lid. Silicone grease was applied on the periphery of the glass chamber coming in contact with the polystyrene material. This type of packaging was done to minimize any possible leakage of sound from inside of the chamber, and also to avoid any interference from external sound. Similar chamber was used to house the ‘control’ (i.e. not subjected to sound stimulation) group test tubes. One speaker was also placed in the glass chamber used for the control tubes at a distance of 15 cm, where no electricity was supplied and no sound was generated. Intensity of sound, measured with a sound level meter (ACD machine control Ltd.) at a distance of 15 cm from the speaker was set in the range of 70-89.5 dB. Sound level in control chamber was found to be below the detection level (40 dB) of the sound level meter. Schematic of the whole experimental set-up is shown in Figure 1. Intermittent mixing of the contents of the test tubes to minimize heterogeneity was achieved by vortexing the tubes at an interval of 3 h using a cyclomixer. Whenever the tubes were taken out for vortexing, each time positions of tubes of a single chamber were inter-changed, and their direction with respect to speaker was changed by rotating them 180°. This was done to achieve a high probability of almost equal sound exposure to all the tubes.

**Figure 1.**
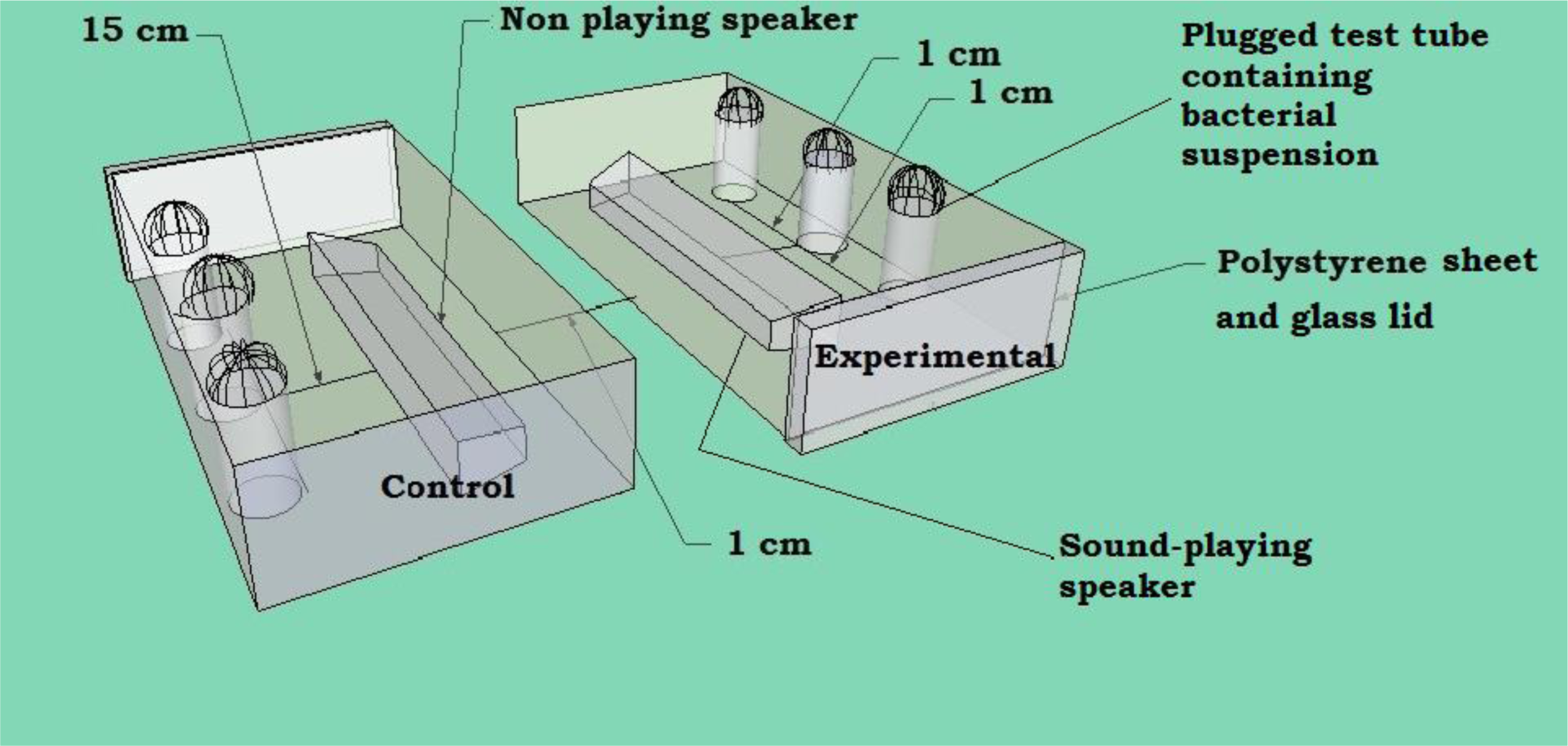
Schematic of the experimental set-up.

### Growth and pigment estimation

#### Violacein extraction

At the end of incubation, after quantifying the cell density at 764 nm (Joshi *et al*., 2016), the *C. violaceum* culture tubes were subjected to extraction of the pigment violacein (Choo *et al*., 2006). Briefly, 1 mL of the culture broth was centrifuged (REMI CPR-24 PLUS) at 12,000 rpm (13,520 g) for 15 min at 25°C, and the resulting supernatant was discarded. The remaining cell pellet was resuspended into 1 mL of DMSO (Merck, Mumbai), and incubated at room temperature for 30 min, followed by centrifugation at 12,000 rpm for 15 min. The violacein extracted in supernatant was estimated by measuring OD at 585 nm. Violacein unit was calculated as the ratio (OD_585_/ OD_764_) indicating violacein production per unit of growth.

#### Prodigiosin extraction (Pradeep *et al.,* 2013)

At the end of incubation, after quantifying the cell density of *S. marcescens* culture at 764 nm, one mL of the culture broth was centrifuged at 10,000 rpm (9,390g) for 10 min. Centrifugation was carried out at 4°C, as prodigiosin is a temperature-sensitive compound. The resulting supernatant was discarded. Remaining cell pellet was resuspended in 1 mL of acidified methanol (4 mL of HCl into 96 mL of methanol; Merck), followed by incubation in dark at room temperature for 30 min. This was followed by centrifugation at 10,000 rpm for 10 min at 4°C. Prodigiosin was obtained in the resulting supernatant; OD was taken at 535 nm. Prodigiosin unit was calculated as the ratio (OD_535_/ OD_764_), which gives an indication of prodigiosin production per unit of growth.

#### Statistical analysis

All the experiments were performed in triplicate, and measurements are reported as mean ± standard deviation (SD). Statistical significance of the data was evaluated by applying *t*-test using Microsoft excel^®^. Data with *p*-values less than 0.05 was considered to be statistically significant.

Additionally, ANOVA was done for all data sets with Microsoft excel^®^. In ANOVA the null hypothesis (that sound treatments at all intensities are equivalent) will not be rejected only if there is no significant difference between any pairs of the means. On other hand, it would be rejected even if there is a significant difference between one pair of means. Therefore it becomes necessary to identify which of the pairs differ significantly and which do not. For this the method given by Snedecor and Cochron in 1959 explained as *Q test* was applied (Gurumani, 2005).

## Results and Discussion

### Effect of speaker with no power supply

Before finalizing the experimental set-up, we investigated whether there is any need to put a non-playing speaker (no power supply, no sound generation) in the control chamber; i.e. whether there is an influence of a silent speaker on the test microorganism. For this particular experiment, the experimental chamber contained a speaker to which no power was supplied, whereas the control chamber contained no speaker. In presence of a speaker gaining no power supply, growth of both the organisms used in this experiment was altered by not more than ~ 3% (Figure 2). Pigment production was affected only in *C.violaceum,* but not in *S. marcescens.* Though the magnitude of alteration in growth and/or pigment formation was not very high, but it was statistically significant. Usually the speakers contain a permanent magnet inside them, besides an electromagnet (http://www.physics.org/article-questions.asp?id=54). The permanent magnet may generate a weak magnetic field, even when electricity supply is not there, which may be responsible for the altered growth and / or pigment production observed in our study. Occurrence of magnetic field effects in biological systems has been mentioned in Evans *et al*., (2013), wherein they have indicated the flavoproteins to be involved in magnetosensitivity of biological entities. Flavoproteins perform a wide range of functions including transcriptional regulation by BLUF proteins in bacteria. Based on the above described observation, we decided to keep a non-playing speaker in the control chamber for all the experiments.

**Figure 2.**
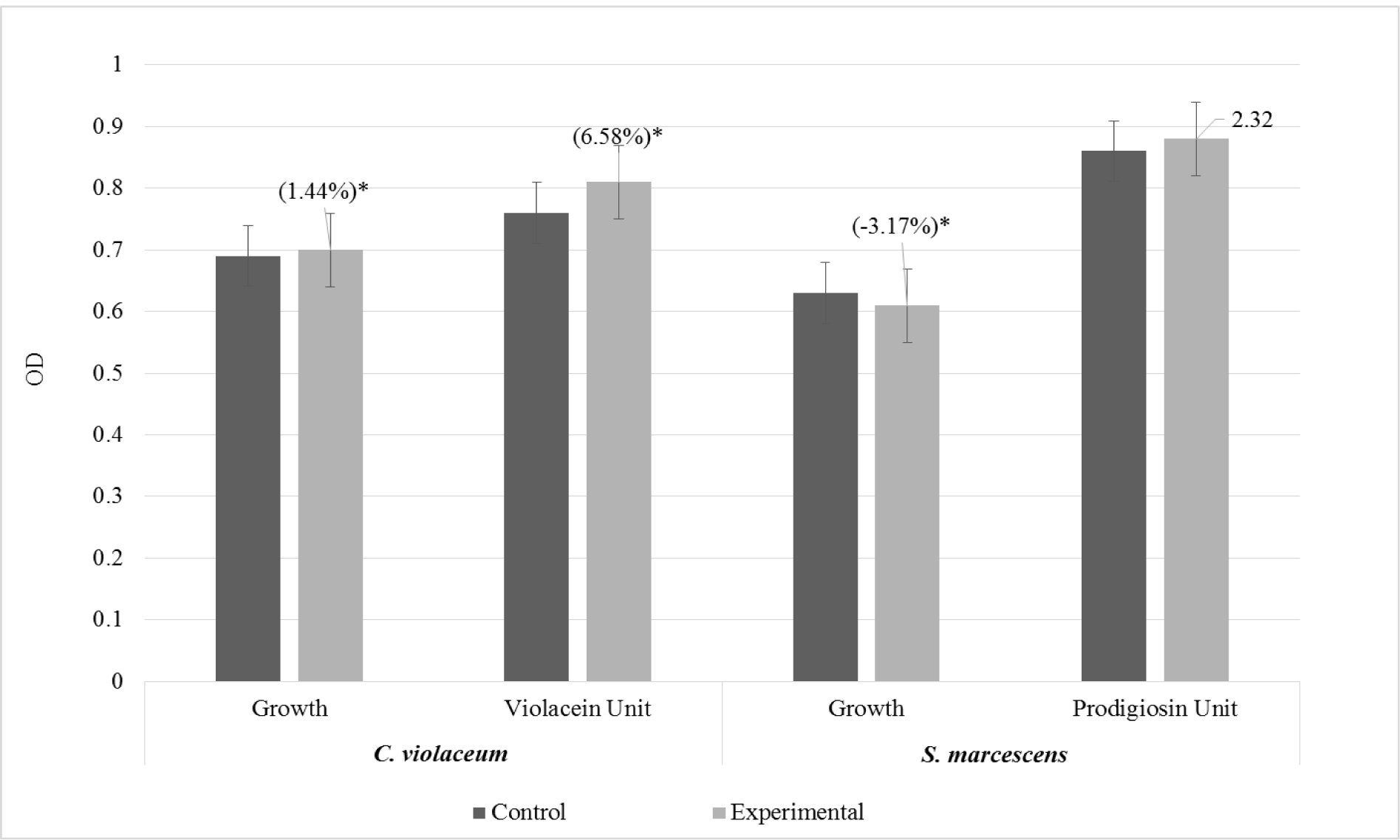
Presence of a speaker with no power supply could influence bacterial growth and/or pigment formation.

### Effect of alteration in sound level at constant frequency

*C. violaceum* culture was exposed to a sound beep corresponding to 300 Hz. Five different levels (each level corresponding to increase of 2 units of the volume regulator of the speaker) of loudness of this sound in the range 70-89.5 dB were tried, keeping the frequency constant. This frequency of 300 Hz was selected based on results of another study (yet unpublished) performed by us, wherein 300 Hz was found to affect quorum sensing (QS) regulated violacein production in *C. violaceum* heavily. Results of the present study (Figure 3) indicated that microbial response to sound at a constant frequency is a function of loudness of the sound, provided there is sufficient change in the intensity of this loudness.

**Figure 3.**
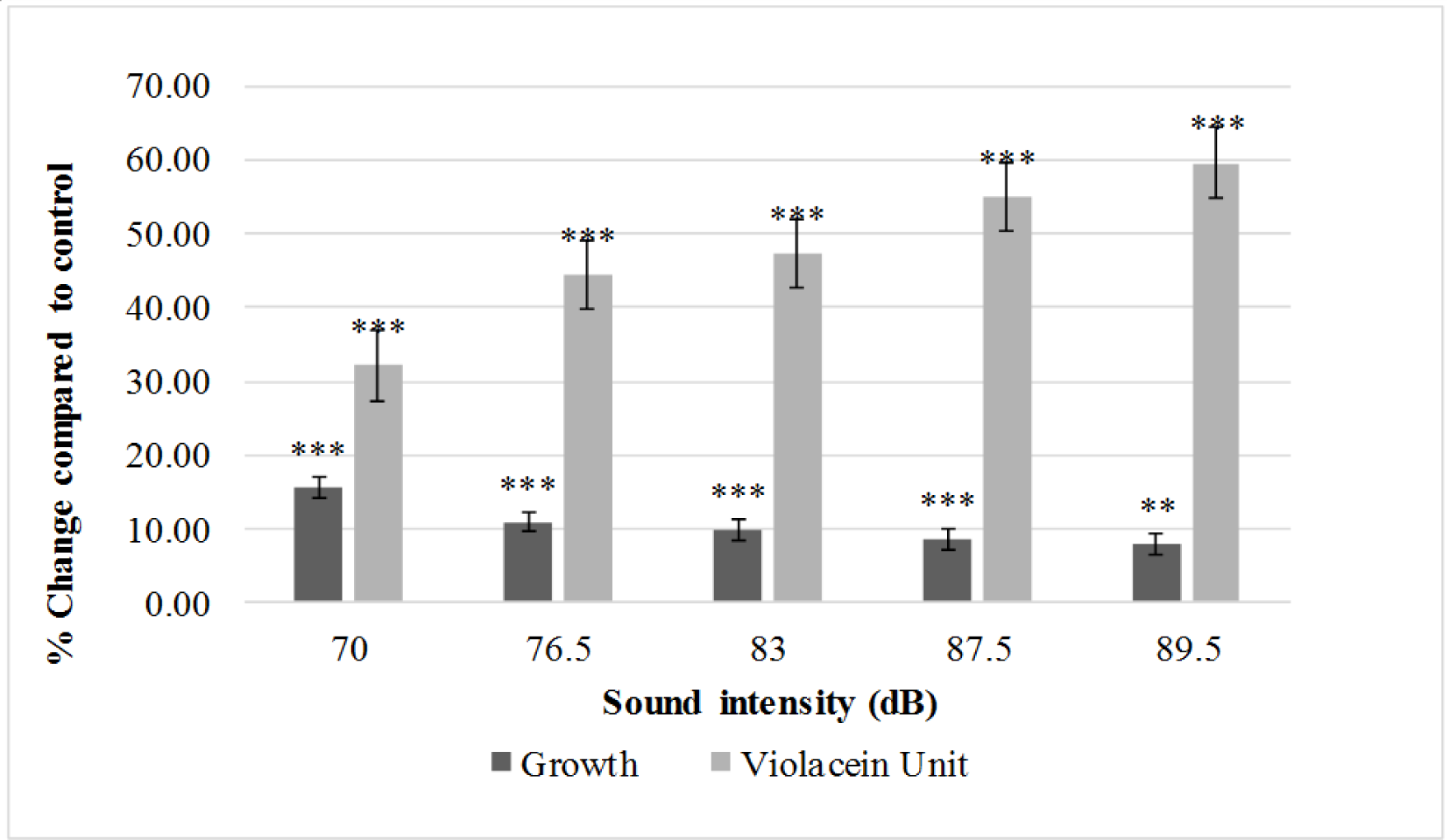
Effect of different levels of sound (300 Hz) intensity on growth and quorum sensing-regulated pigment (violacein) production in *C. violaceum*.

Growth of *C. violaceum* was affected the most (15.02%) at the lowest level (70 dB) of sound intensity tested, whereas its QS-regulated pigment production was affected the most (59.63%) at the highest level (89.5 dB) of sound intensity tested. Effect of sound stimulation on growth of *C. violaceum* was observed to decrease with increase in sound intensity. On the contrary, effect of sound stimulation on pigment production was observed to increase along with increase in sound intensity. With respect to growth, the *Q test* indicated that for any two values of ‘percent change’ in bacterial growth to be considered significantly different, the minimum difference between them has to be 3.57. We found that to achieve this critical difference of 3.57%, the difference in the loudness level has to be >13 dB. For example, the effect of 300 Hz at 70 dB on *C. violaceum* was statistically not different than that at 76.5 dB as the difference (6.5 dB) between these two loudness levels was < 13 dB. On the other hand, the effect of 70 dB sound was significantly different from that on 83 dB and above, as here the difference between the concerned sound levels exceeded 13 dB. Similarly with respect to violacein unit, the significant difference (14.29%) corresponded to a decibel difference of 13 again. These results suggest that loudness of the sound (of an appropriate frequency) can be a significant factor causing alteration in bacterial growth and production of QS-regulated production of pigment, provided that the change in the level of loudness exceeds a certain critical level. Gu et al. (2013) tried different combinations of sonic frequency and intensity with *Escherichia coli*, and reported that irrespective of what combination is tried, growth, protein content and enzyme activity (catalase and SOD) of *E. coli* experienced an upward effect owing to sonic treatment.

### Effect of the size of test tube in which bacteria are exposed to sound stimulation

To investigate whether size of the vessel in which bacteria are exposed to sound can make any difference, *C. violaceum* was exposed to sound of five different frequencies (100 Hz, 200 Hz, 300 Hz, 500 Hz, and 1,000 Hz) in test tubes of two different dimensions. Sound level (measured in dB; Table 1) and media volume (6 mL) was kept same for both sizes of the test tube. Results of this experiment presented in Figure 4, indicate that size of the vessel in which bacteria are exposed to sonic stimulation did not make a difference with respect to their response to the external sound stimuli, in majority of cases. For example, with respect to cell density, size of the test vessel made significant different only at 500 Hz; whereas this size factor had no significant impact on the pigment production at any of the five test frequencies. However, such studies needs to be conducted in still bigger vessels too, as the sound power (energy per unit area) may vary more in case of larger vessels. Kram and Finkel (2014) had shown culture volume and vessel to affect long-term survival, mutation frequency, and oxidative stress of *Escherichia coli*, and they suggest that culture vessel and incubation conditions should be carefully considered in the planning or analysis of experiments. However, Hammes *et al.*, (2010) while analysing the impact of surface-to-volume ratio on final bacterial concentrations after batch growth, examined six bottle sizes (20 to 1,000 ml) using three independent enumeration methods to quantify growth, and found no evidence of a ‘volumetric bottle effect’. Thus, one come across contradictory findings in literature concerning the effect of vessel size. In our present study too, results at 500 Hz indicate vessel size to make a difference to bacteria growth, but not so at other test frequencies.

**Table 1.** Level of sound intensity at different frequency.

| **Sr. No** | **Frequency of the test sound (Hz)** | **#Intensity as measured by the sound level meter (dB)** |
| --- | --- | --- |
| 1 | 100 | 80 |
| 2 | 200 | 85 |
| 3 | 300 | 90 |
| 4 | 500 | 84 |
| 5 | 1000 | 86 |

<sup>#</sup> Speakers were set at a constant level of 9 (volume indicating unit) for all frequencies.

**Figure 4.**
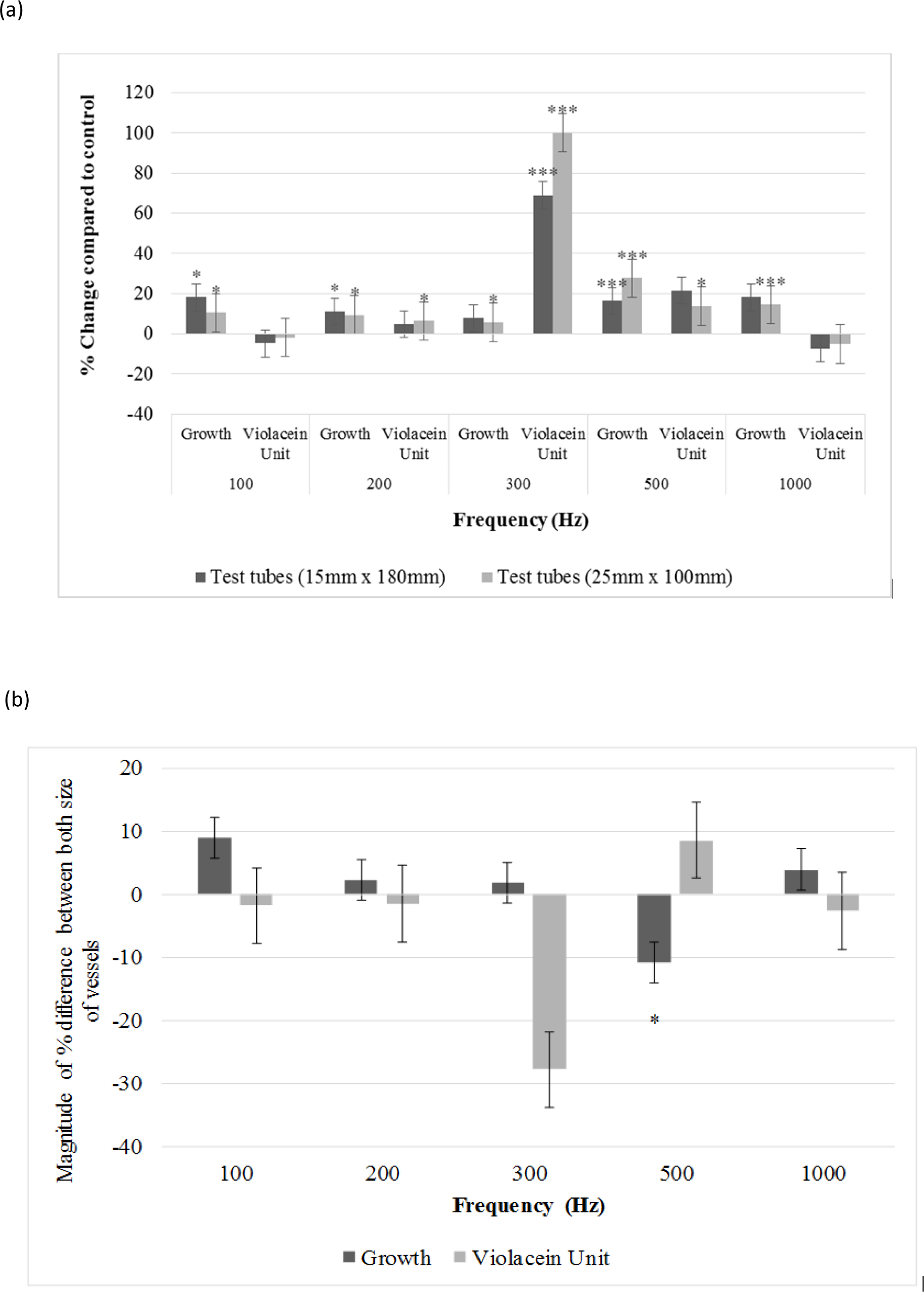
(a) Effect of different frequencies of sound on growth and pigment production of *C. violaceum* in two different size of test tubes; (b) Magnitude of difference in ‘percent change’ values for both (growth and pigment production) parameters between two vessel sizes.

## Conclusion

In one of our previous unpublished study, we found that the quorum sensing-regulated production of the pigment violacein in *C. violaceum* is affected heavily by providing sonic stimulation of 300 Hz to this bacterium. In the present study, we exposed this bacterium to 300 Hz stimulation at five different levels of intensity in the range of 70-89.5 dB, for investigating whether bacterial response to sonic stimulation can be a function of the sound-level. Our results indicated that magnitude of the bacterial response to sound exposure can change notably depending on the level of sound intensity, provided the difference between the sound-levels tested exceeds the minimum critical value, which in this case was found to be 13 dB. Growth of *C. violaceum* was affected more at lower sound intensity, whereas pigment production was affected more at higher intensities, suggesting that the quorum sensing machinery of this bacterium responds differently to external sound stimuli than the growth related cellular machinery. Further in this study, we also investigated whether a silent speaker emitting no sound can exert any effect on the test bacteria (*C. violaceum* and *S. marcescens*). Silent speaker was indeed found to affect growth of both, and pigment production in *C. violaceum* to a smaller (but statistically significant) extent, suggesting that any studies regarding effect of sound on microbes should include non-playing speaker as ‘control’. This may be due to the ability of some of the cellular components like flavoproteins to respond to the external magnetic field effects. We also investigated whether size of the test-tube in which bacteria were exposed to sound can make any difference, however vessel size was not found to be a major factor influencing the experimental outcome. In this study, besides measuring cell-density, we also measured pigment production of the test cultures. It should be noted that pigment production is just one of the many parameters in these bacteria which is regulated by quorum-sensing. Hence, it is possible that in fact many other quorum sensing associated parameters in bacteria may be affected by external sonic and/or magnetic stimuli. Based on our primary findings, we decided to compare the gene-expression (whole transcriptome) profile of the 300 Hz sound-treated *C. violaceum* culture with its counterpart receiving no sonic stimulation. Currently, we are analysing results of the said experiment, and shall be reporting the same in a separate paper in near future.

## Acknowledgement

Authors thank Nirma Education & Research Foundation (NERF), Ahmedabad for financial and infrastructural support; Vidhi Shah and Jinal Sukhadiya for help in manuscript formatting.

